# Deep learning-based method to identify disease-resistance proteins in *Oryza sativa* and relative species

**DOI:** 10.1101/2023.11.25.568625

**Authors:** Vedikaa Dhiman, Soham Biswas, Asish Kumar Swain, Ayan Sadhukhan, Pankaj Yadav

## Abstract

Rice (*Oryza sativa*) is a significant agricultural crop consumed by more than half of the global population. Its demand is expected to increase due to rising consumption and a growing global population. Moreover, the rice plant is frequently exposed to disease-causing pathogens, such as bacteria, fungi, viruses, and nematodes. Thus, cultivating disease-resistant varieties is an efficient way of disease control compared to pesticide applications. However, the rice plant has a well-defined defense system to prevent the onset of disease, including Pathogen-associated molecular pattern (PAMP)-triggered immunity (PTI) and effector-triggered immunity (ETI). The defense system is controlled by various disease-resistance proteins, such as resistance (R) proteins and pathogen recognition receptors (PRRs). Therefore, the identification of disease-resistance proteins not only reduces the amount of pesticides used in rice fields but also increases their yield. Though some resistant proteins have been characterized, their rapid identification, precise diagnosis, and appropriate management are still lacking. However, few methods based on sequence-similarity and de novo prediction, such as Machine Learning (ML), usually have low prediction power. In this study, we built a state-of-the-art classifier based on Deep Learning (DL) for the early detection of disease-resistance proteins in rice and related species. We compared the DL-based Multi-layer Perceptron (MLP) model with the five well-established ML-based methods using a protein dataset of rice and its related species. The DL-based MLP model outperformed all of the five classifiers on 10-fold cross-validation. The accuracy, Area Under Receiving Operating Characteristic (ROC) curve (AUC), F1-score, precision, and recall were superior in the DL-based MLP model. In conclusion, the MLP model is an effective DL model for predicting disease-resistance proteins with high scores in performance metrics. This study will provide insight to the breeders in developing disease-resistant rice varieties and assist in transforming traditional rice farming practices into a new age of smart rice farming.

## INTRODUCTION

Rice (*Oryza sativa*) is the primary staple food crop consumed by more than half of the world’s population, and it is a model monocot species for plant research. A wide range of biotic stresses can significantly reduce rice yield and quality. These stresses include pathogenic bacteria, fungi, viruses, nematodes, pest attacks, and invasion by parasitic plants. Moreover, biotic stresses have been identified as major contributors to reducing rice yield, with a global estimate of approximately 30.0% in 2019 (Vo et al., 2021). Currently, the primary methods of combating various diseases are using pesticides and deploying disease-resistant cultivars. However, excessive use of pesticides not only increases the cost of rice production but also causes considerable environmental damage. However, fewer translational approaches have explored the rice-inducible defence mechanisms, including PTI and ETI. The PTI is governed by PRRs, resulting in the avirulence (Avr) pathogen effector recognition. In contrast, cytoplasmic immunoreceptors control ETI by activating the R genes (Kumar et al., 2022). Recently, a cognate resistance gene, *Xa7*, has been identified as displaying durable resistance in rice varieties under field conditions (Zhao et al., 2022). In another study, a novel nucleotide-binding domain and leucine-rich repeat-containing protein (NLR) family *Xa47* gene have been identified with broad-spectrum resistance to bacterial blight disease (Lu et al., 2022). Furthermore, a recent proteomic study indicated that infected transgenic rice plants exhibit stimulation in the protein 14–3–3GF14f, which is involved in signal transduction pathways (Molla et al., 2020).

Experimental methods are the traditional ways to find disease-resistance proteins in rice. Early detection of these proteins facilitates the process of genetic improvement and breeding programs for developing disease-resistant varieties (Pal et al., 2016). However, these methods are time-consuming and known to have a high false positive rate. Thus, new efficient and accurate methods are required, considering the importance of the role of disease-resistance proteins. In this regard, efforts have been made in the past years to detect disease-resistance proteins rapidly with high precision through various computational methods. In the last few years, ML and DL approaches have been extensively used to detect disease-resistance proteins in rice and other related species. Recently, several ML-based approaches have been used, from disease classification to disease-resistance protein identification. In a current study, a support vector machine (SVM) based approach was used to classify the biotic or abiotic stress condition using microarray technology having 559 samples and 13 stress situations (Das et al., 2022). Similarly, RLPredictOme correctly predicts Leucine Rich Repeats-receptor-like proteins (LRR-RLPs) that stimulate an immune response against disease in rice (Silva et al., 2022). Moreover, ML methods are easier to apply and do not need a huge amount of training data; however, such techniques are time-consuming and need human expertise. Further, ML-based algorithms consist of computational complexity and detection robustness.

Recently, DL methods such as convolutional neural networks (CNNs) and deep belief networks (DBNs) were found to detect plant diseases accurately (Shoaib et al., 2023). Similarly, prPred-DRLF, a DL-based model, was developed to predict plant resistance (R) protein (Wang et al., 2022). In addition, several DL-based approaches were developed based on rice leaf images for disease classification. A DL-based faster region-based convolutional neural network (Faster R-CNN) was developed to diagnose rice leaf diseases (Bari et al., 2021). Further, a DL and SVM-based model was developed to recognize four rice leaf diseases through their images (Jiang et al., 2020). However, various image-based DL models are available to detect diseases, but none of them exploit the protein sequence information of the rice.

To address these challenges, we developed a deep neural network-based approach for disease-resistance protein identification in *Oryza sativa* and relative species. Our approach requires the amino-acid sequence of the query protein to identify the disease resistance status. First, major features were extracted for the protein sequence using several approaches, and then, a DL model was employed that predicted the disease resistance level with greater accuracy. We utilised a state-of-the-art approach to build the DL model by integrating proteomics data with ML techniques. We retrieved known disease resistance and a whole set of proteins from 13 plant species related to *Oryza sativa*. Important features were then extracted and validated on a broad set of proteins in related rice species. Next, several ML and DL-based approaches were tested on the extracted datasets. Further, all models were tested and validated using 10-fold cross-validation. Our developed approach is publicly available for protein disease-resistant classification on GitHub.

## MATERIALS & METHODS

This section describes the computational framework of the DL model developed to predict disease-resistance proteins in rice and related species. The framework includes data retrieval, data filtering, feature extraction, selection methods, and application of ML and DL methods. A pipeline of the proposed methodology is depicted in **Figure 1**, defining the ordered set of steps necessary for its execution.

**Fig. 1.**
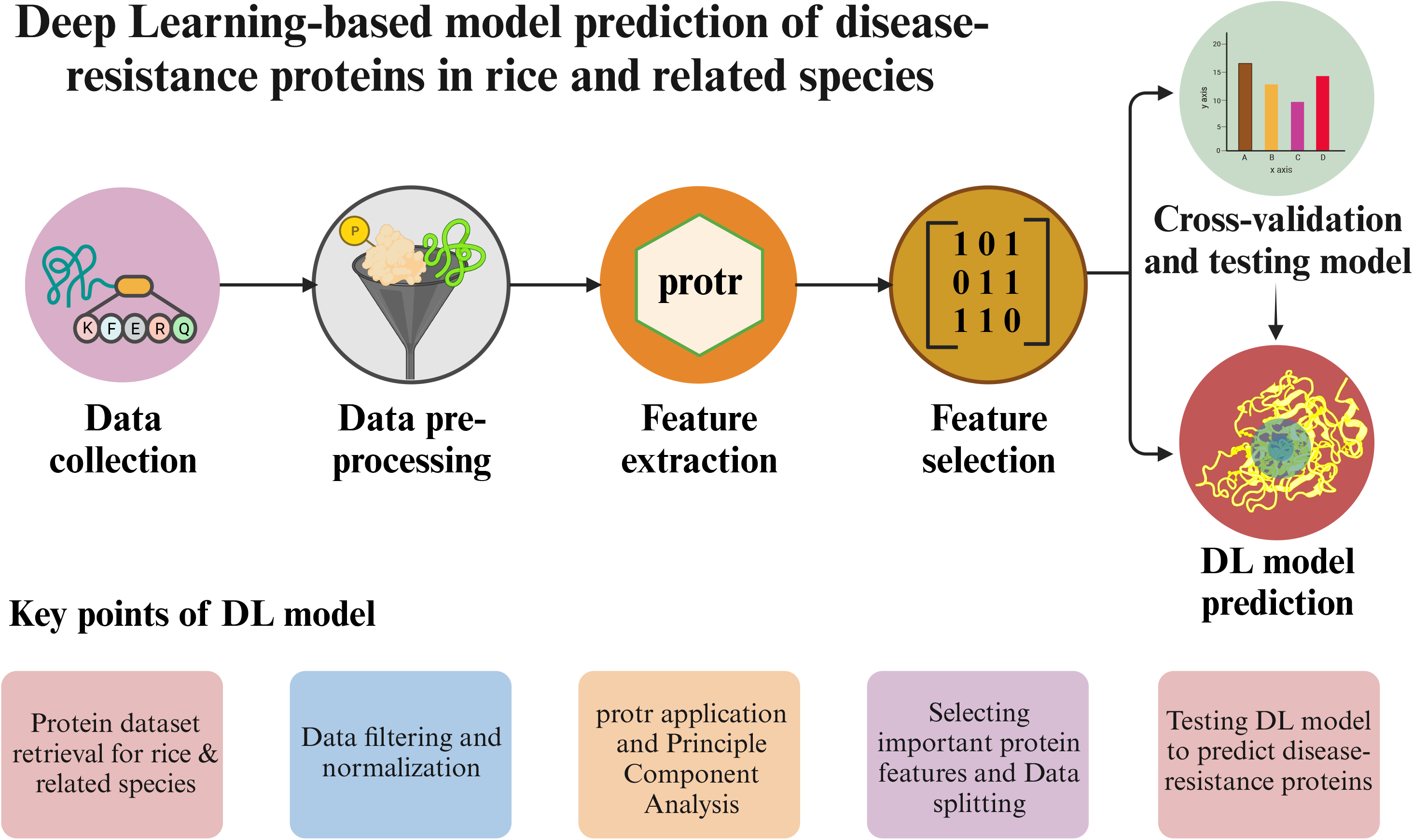
Shows a pipeline of the proposed methodology.

### Data collection

A collection of 98 experimentally validated disease-resistance proteins from 13 plant species closely related to *Oryza sativa* have been retrieved from the RiceRelativesGD V4.0 database (Mao et al., 2019). These plant species include *Oryza nivara, Oryza rufipogon, Oryza glaberrima, Oryza barthii, Oryza glumaepatula, Oryza meridionalis, Oryza punctaca, Oryza brachyantha, Zizania latifolia, Echinochloa crus-galli, Oryza sativa* f. *spontanea* (*indica* type), *Leersia perrieri*, and *Oryza sativa* subsp. *indica* group. These 98 proteins extracted from the RiceRelativesGD V4.0 database served as a positive dataset. For the negative dataset, the entire proteome from the same 13 species was also obtained from the RiceRelativesGD V4.0 database. We have extracted 6,51,535 proteins as a negative dataset. **Table S1** shows details of databases and tools used in the study.

### Data preprocessing

The retrieved dataset has a significant data imbalance, with only 98 proteins in the positive dataset and a substantial 6,51,535 proteins in the negative dataset. To handle the data imbalance, previous studies randomly selected negative datasets from the whole proteome (Pal et al., 2016). However, this approach can introduce biases into the model, as unannotated disease-resistance proteins may be present in the proteome. Additionally, the model will heavily depend on selecting negative criteria randomly. To address these challenges, we employed three criteria to balance the datasets instead of randomly selecting protein sequences from the negative dataset.

Firstly, we removed proteins with disease-resistant domains such as Leucine Rich Repeats (LRR), Kinase, Nucleotide Binding sites (NBS), and other similar domains from the negative dataset. These disease-resistant domains were identified by utilizing the Disease Resistance Analysis and Gene Orthology (DRAGO 3) tool of the Plant Resistance Genes Database (PRGdb) (Calle García et al., 2022). Secondly, protein sequences containing incomplete fragments, i.e., ‘X’ sequences, were removed. These ‘X’ sequences are found to be unknown and were present in between the protein sequences. Thirdly, the NCBI-BLASTP (Johnson et al., 2008) was performed to reduce the size of the negative dataset. The BLASTP was conducted based on percent identity (>95%), query coverage (>75%), and e-value (1e-6) for the protein sequences of the negative dataset. In addition, the data redundancy was further reduced by using the Cluster Database at High Identity with Tolerance (CD-HIT) toolkit (Fu et al., 2012). It is a program for clustering biological sequences to reduce sequence redundancy and improve the performance of other sequence analyses.

### Feature extraction and selection

After balancing the positive and negative datasets, we extracted and selected important features for constructing the model. Thus, the protr package (Xiao et al., 2015) in the R program was used to extract important features of the protein sequences. The protr package involves the generation of different numerical representation schemes of proteins and peptides from amino acid sequences. It includes various important features or descriptors such as amino acid composition, autocorrelation, CTD (composition, transition, distribution), conjoint triad, etc. The details of the descriptors are described below:

#### 1. Amino acid composition (AAC)

It describes the fraction of each amino acid type within a protein sequence and can be shown as follows:

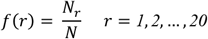

Here *N*_*r*_ is the number of the amino acid type r, and N is the sequence length. In this study, the sequence length was the same as that of positive and negative datasets, i.e., 98 and 198, respectively.

#### 2 Autocorrelation

It describes amino acid properties along the sequence and is implicated by:

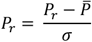

Here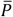 is the average of the property of the twenty amino acids:

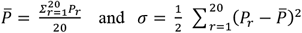

#### 3 Composition, Transition, Distribution (CTD)

Composition (C) describes the overall distribution of amino acids in a protein sequence, transition (T) describes the frequency with which different amino acids are adjacent to each other, and distribution (D) describes the distribution of amino acids along the length of the protein sequence. The composition is defined as:

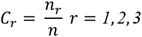

Here *n*_*r*_ is the number of amino acid type r in the encoded sequence; n is the sequence length. Transition is defined as:

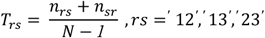

Here n_rs_ and n_sr_ are the numbers of dipeptide encoded as “rs” and “sr” in the sequence, and N is the sequence length.

#### 4 Conjoint Triad Descriptors

CTD classifies the amino acids into seven groups, representing the frequency of each triad (a sequence of three amino acids) in a protein sequence. CTD is used to identify patterns in protein sequences that are associated with protein-protein interactions.

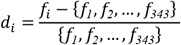

The numerical value of d_i_ of each protein ranges from 0 to 1, enabling the comparison between proteins, and f_i_ is the i^th^ value of frequency of the triads of the protein sequences.

#### 5 Quasi-sequence-order descriptors

The quasi-sequence-order descriptor is derived from the distance matrix between twenty amino acids measures. For a particular amino acid, the quasi-sequence-order is calculated by adding the distances between one amino acid and all other amino acids in the protein sequence. The distance matrix between the twenty amino acids is calculated by:

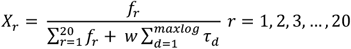

Here, f_r_ is the normalized occurrence for amino acid type i, and *w* is a weighting factor (*w* = 0.1). The *d*-th rank sequence-order-coupling number is defined as:

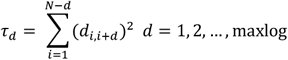

where, d_i, i+d_ is the distance between the two amino acids at position i and i+d.

#### 6 Pseudo-amino acid composition (PseAAC)

The PseAAC of a protein includes a set of discrete numbers derived from its amino acid sequence and different from the classical amino acid composition. It is based on classifying amino acids into twenty groups, and it represents the frequency of each group in the protein sequence. PseAAC is generated by 20+λ discrete numbers to represent a protein and can be defined as:

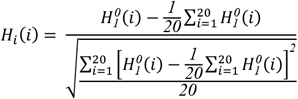

Where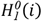is the original hydrophobicity value.

#### 7 Profile-based and Proteochemometric modeling descriptors

The feature vectors of profile-based methods are based on the Position-Specific Scoring Matrix (PSSM) by running PSI-BLAST and showing good performance. On the other hand, proteochemometric modeling includes statistical modeling techniques to model ligand-target interaction space.

**Table S2** depicts a list of features of the protr package used in the study.

#### 8 Feature selection

The features extracted using the protr package yielded large-sized matrices for each linear and non-linear descriptor. Therefore, feature selection was carried out from the pool of descriptors to reduce the descriptor vector dimensions and include only relevant descriptors that will generate meaningful data. This was necessary to minimize computational time and complexity.

The matrices generated by feature extraction were examined manually. Few descriptors resulted in null outputs for a certain feature vector calculation. After conducting manual tests on whether the null outputs are overwhelming among feature vector components, the respective descriptor related to that feature vector was discarded. The f-score, related to the feature characteristics, was then calculated for all the descriptors and compared against the already benchmarked control dataset given by the F-select protocol of the libSVM (Chang, et al., 2007). Finally, features were ranked out based on the calculated f-scores and aligned their functional descriptions with the objective of the study. Further, based on the feature select test results, we selected all the above-mentioned descriptors from AAC to PseAAC. Among profile-based descriptors, extractPSSM and extractPSSFeature were selected. In addition, from a set of proteochem[ometric modeling descriptors, BLOSUM (AABLOSUM62) and PAM (AAPAM120) matrix-based scales, amino acid properties-based scales (AAindex), molecular topology-based scales (AAMOE2D, AAMOE3D, AATopo, AAGeom, AAInfo, AAmolProp) and weighted holistic invariant molecular (WHIM) based descriptor scale AAWHIM were selected. The remaining seventeen descriptors from this set were discarded to build the model.

### Modeling of disease-resistance protein classifiers

The positive and pre-processed negative datasets were split into 80% as training and 20% as test sets using *Scikit-learn* (Pedregosa FABIANPEDREGOSA et al., 2011) in the Python program. This package includes various advanced ML and DL algorithms for supervised and unsupervised problems. In addition, as the dataset has 19503 features, principal component analysis (PCA) was performed to reduce the dimension (“Advances in Neural Information Processing Systems 10 - Proceedings of the 1997 Conference, NIPS 1997,” 1998) PCA applied with the number of components set at 50 covers the 61.09% variance of the dataset.

We have used both statistical, i.e., logistic regression (LR), ML, and DL classifiers to predict disease-resistance proteins. These classifiers include Random Forest (RF), Support Vector Machine (SVM), Naïve Bayes (NB), K-Nearest Neighbor (KNN), and MLP. The parameters described below for each ML method used for the model building are:

#### 1. Logistic Regression

It is the statistical algorithm for classifying categorical class variables (Kirola et al., 2022). It analyses the relationship between a set of independent variables and dependent binary variables. It is based on the desire to model the probabilities of the output classes while ensuring that output probabilities sum to one and remain between zero and one, as expected from probabilities.

#### 2. Random Forest

It is a collection of predictable trees in which each tree is based on independently sampled random vector values with the same distribution of all the trees in the forest (Chauhan et al., 2021). It consists of a collection of tree-structured classifiers {h (x, Θ_k_), k = 1, …} where the {Θ_k_} is independent identically distributed random vectors, and each tree casts a unit vote for the most popular class at input x (Breiman, 2001). While performing RF, the parameters for the model include 100 as ‘n_estimators’ and 0.01 as ‘ccp_alpha.’

#### 3. Support Vector Machine

It is a technique for supervised ML that can analyze data and recognize patterns and is extensively used for classification and regression analysis (Pal et al., 2016). It is based on the decision planes that define decision boundaries. It includes nonlinear functions known as kernels to transform input space into a multidimensional space.

#### 4. Naïve Bayes

It is a probabilistic classification method that employs the Bayes theorem and determines the probability of each feature occurring in each class within a dataset using a set of features from the dataset (Morgan et al., 2021). The values of the attributes are used to calculate the posterior probability for each class within the dataset for each row of data. The row of data is then assigned to the class with the highest posterior probability.

#### 5. K-Nearest Neighbor

KNN performs classification and prediction based on locating the K nearest matches in training data and then using the label of the closest matches (Kartikeyan & Shrivastava, 2021). It extracts the most similar k data points from the training set and assigns the new data points to the category where K similar data points occur most frequently. The value of K was considered to be one for the model classification.

#### 6 Deep learning

MLP is a non-linear computational method used for the classification and regression of complex systems (Yoosefzadeh-Najafabadi et al., 2021). This algorithm includes several highly interconnected processing neurons that can be used in parallel to detect a solution for a specific problem. For our analysis, we have used ‘sgd’ as a solver with 1000 as hidden layers. In addition, epoch as 1000, activation function as ‘relu,’ and learning rate as 0.001 were considered.

A confusion matrix was constructed for each model using the previously created test dataset to validate the models. The confusion matrix of the models includes information regarding true positive (TP), false positive (FP), true positive (TP), and true negative (TN). Additionally, various metrics such as accuracy, AUC, F1-score, precision, and recall were calculated through 10-fold cross-validation. Detailed descriptions of these metrics are provided below:

#### 1. Accuracy

It is the percentage of correctly classified data points from the total number of samples (Khalili et al., 2020) and is described as:

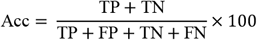

#### 2. ROC curve

It is used to summarize the performance of a binary classifier in a single number.

#### 3. F1-score

It shows the harmonic mean of recall and precision and can be calculated as follows (Khalili et al., 2020):

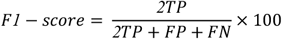

#### 4. Precision

It is also known as positive prediction value (PPV), which compares the number of correctly classified positive samples to the total number of positive samples predicted (Khalili et al., 2020). It can be explained as:

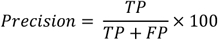

#### 5. Recall

It measures the ability of the model to detect positive samples. The higher the recall, the more positive samples detected. It can be shown as:

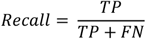

## RESULTS

### Optimization of datasets

The study took into account disease-resistance proteins as positive and whole proteome as negative datasets of 13 closely related species of *Oryza sativa*. A total of 98 experimentally validated disease-resistance proteins were considered as positive datasets, whereas the negative dataset consists of 6,51,535 proteins. In the negative dataset, 3,33,342 proteins consisting of disease-resistant domains were identified and removed. We identified six types of disease-resistant domains in the negative dataset. These domains were LRR, Kinase, Transmembrane (TM), receptor-like kinase (RLK), receptor-like protein (RLP), coiled-coil (CC), Toll-interleukin-1 receptor (TIR), and NBS. Thus, after the removal of proteins with disease-resistant domains, we found a total of 3,18,193 proteins. In addition, we have found and excluded 9007 ‘X’ sequences in these protein sequences, and 3,09,186 proteins were finally obtained. These ‘X’ sequences were found to be unknown sequences or act as a stop codon in the protein sequence. Next, NCBI-BLASTP was performed on these proteins to normalize both datasets further. It was based on an e-value of 1e-6, query coverage of>75%, and percent identity of >95%, and a total of 45,379 proteins were extracted. Furthermore, clustering by CD-HIT was performed to reduce the redundancy among the datasets. In the protein sequences of the negative dataset, 33 clusters were identified, each consisting of one representative sequence. These 33 clusters in the negative dataset were identified based on two parameters. The parameters used include the word size of protein sequences as five and the threshold for clustering as 0.7∼1.0. Among these clusters, a total of nine clusters with sequences having 95% to 100% similarity with the representative sequences were considered for further analysis. Consequently, we obtained 98 proteins as a positive dataset and 198 proteins as a negative dataset for feature extraction and selection.

### Feature analysis of models

Both datasets were balanced, and 19,503 features were extracted using the protr package for each protein sequence. These features represent the structural, functional, expression, and interaction profiles of the proteins. A dimension reduction was performed with the top fifty principal components to reduce the feature dimension. In addition, six ML and DL algorithms were employed to assess the protein disease-resistance classification performance. These algorithms include LR, RF, SVM, NB, KNN, and MLP.

The positive and negative datasets were split into train and test sets. The splitting was performed to eliminate biases so that both the train and test sets have an equal proportion of positive and negative datasets. **Table 1** summarizes the number of proteins in the training and test sets. The confusion matrix generated from all the models is shown in **Figure 2**. All ML methods, excluding the SVM, performed well on the test set. Detailed information regarding TP, FP, TN, and FN for the models is described in **Table 2**.

**Table 1.** Division of positive and negative datasets into training and test sets.

| Datasets | Training set | Test set |
| --- | --- | --- |
| Positive | 78 | 20 |
| Negative | 157 | 41 |

Datasets: positive and negative datasets having disease-resistance protein and whole proteome set, respectively; Training set: a subset to train the model; Test set: a subset to test the trained model.

**Table 2.** TP, FP, TN, and FN values of the models.

| Classifiers | TP | TN | FP | FN |
| --- | --- | --- | --- | --- |
| LR | 37 | 18 | 3 | 2 |
| RF | 40 | 18 | 3 | 2 |
| SVM | 38 | 17 | 2 | 3 |
| NB | 38 | 17 | 2 | 3 |
| KNN | 36 | 18 | 4 | 2 |
| MLP | 65 | 16 | 1 | 4 |

Classifiers: name of all six classifiers, i.e., Logistic Regression (LR), Random Forest (RF), Support Vector Machine (SVM), Naïve Bayes (NB), K-Nearest Neighbor (KNN) and Multilayer perceptron (MLP); TP: true positives is the number of times our actual positive values are equal to the predicted positive; TN: true negatives is the number of times our actual negative values are equal to predicted negative values; FP: number of times our model wrongly predicts negative values as positives; FN: false negative is the number of times our model wrongly predicts negative values as positives.

**Fig. 2.**
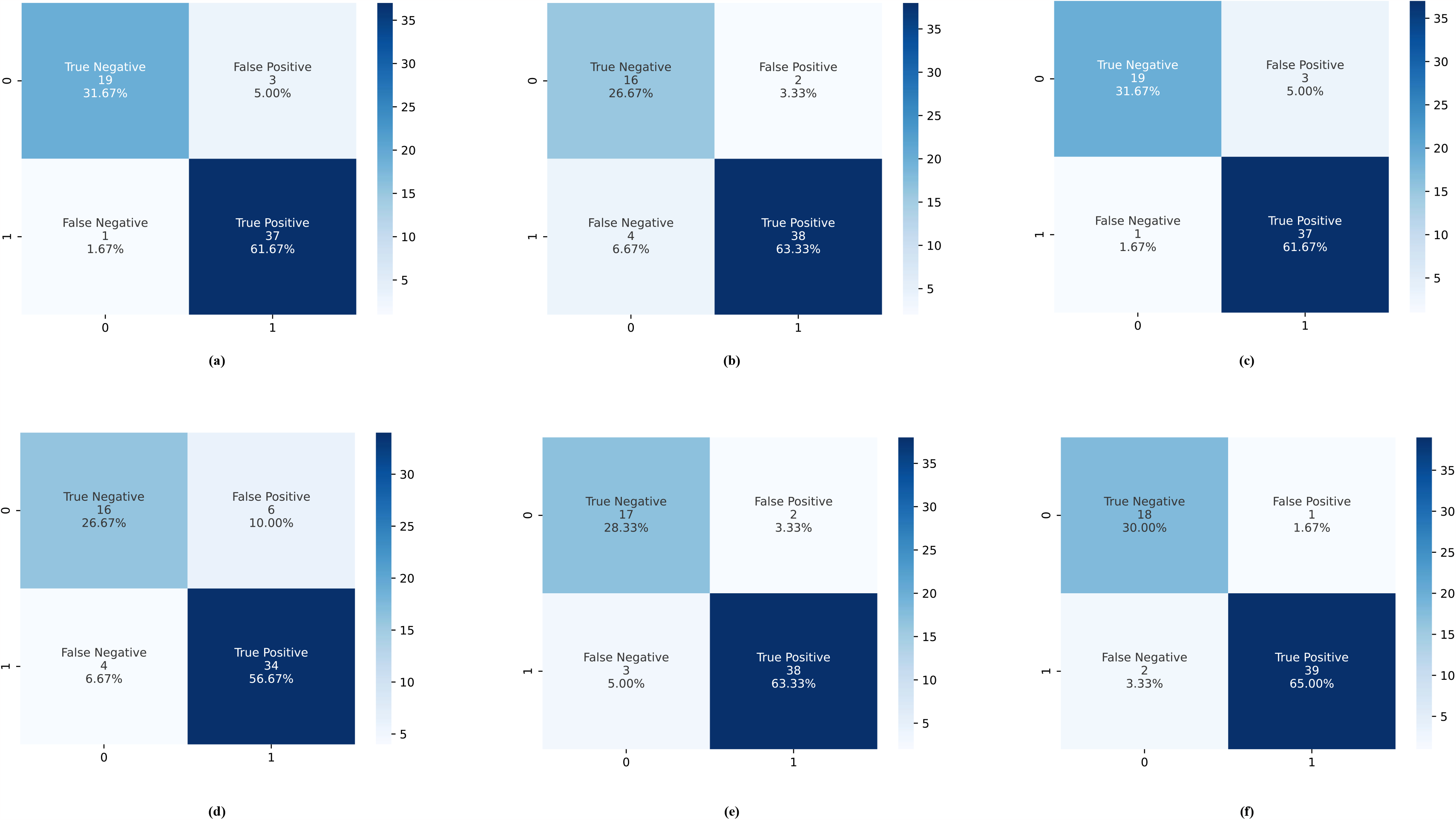
Shows the confusion matrix performance measures of the models. (a-f) shows a confusion matrix of LR, RF, SVM, NB, KNN, and MLP, respectively.

### Model evaluation

Evaluation of the model is crucial in order to check the overfitting of the model on the test sets. Hence, the performance of the models predicting disease-resistance proteins was evaluated by the 10-fold cross-validation method. This method is a commonly used model validation technique where a subset of data is preserved by testing once or repeatedly. Thus, various parameters, including accuracy, AUC, F1-score, precision, and recall, were calculated for the six classifiers. Among all the six classifiers, the DL-based MLP model showed the highest mean validation accuracy of 91.59% with a training accuracy of 96.39%. In addition, MLP showed the best performance, having accuracy, AUC, F1-score, precision, and recall of 91.59%, 97.84%, 93.77%, 95.65%, and 92.5, respectively. **Figure 3** highlights a comparative analysis of the parameters of the models. **Table 3** illustrates the performance evaluation of the models.

**Table 3.** Performance evaluation of the models.

| Models | Performance metrics |  |  |  |  |
| --- | --- | --- | --- | --- | --- |
|  | Accuracy | AUC | F1-score | Precision | Recall |
| LR | 84.17 | 91.05 | 86.47 | 92.39 | 81.86 |
| RF | 88.89 | 95.31 | 91.39 | 91.24 | 92.00 |
| SVM | 79.06 | 87.36 | 84.80 | 90.68 | 80.39 |
| NB | 85.52 | 92.96 | 86.84 | 95.37 | 80.31 |
| KNN | 88.91 | 88 | 91.96 | 92.47 | 92.00 |
| MLP | 91.59 | 97.84 | 93.77 | 95.65 | 92.50 |

Models: name of the models; Performance metrics: performance parameters of the models, including accuracy, AUC, F1-score, precision and recall.

**Fig. 3.**
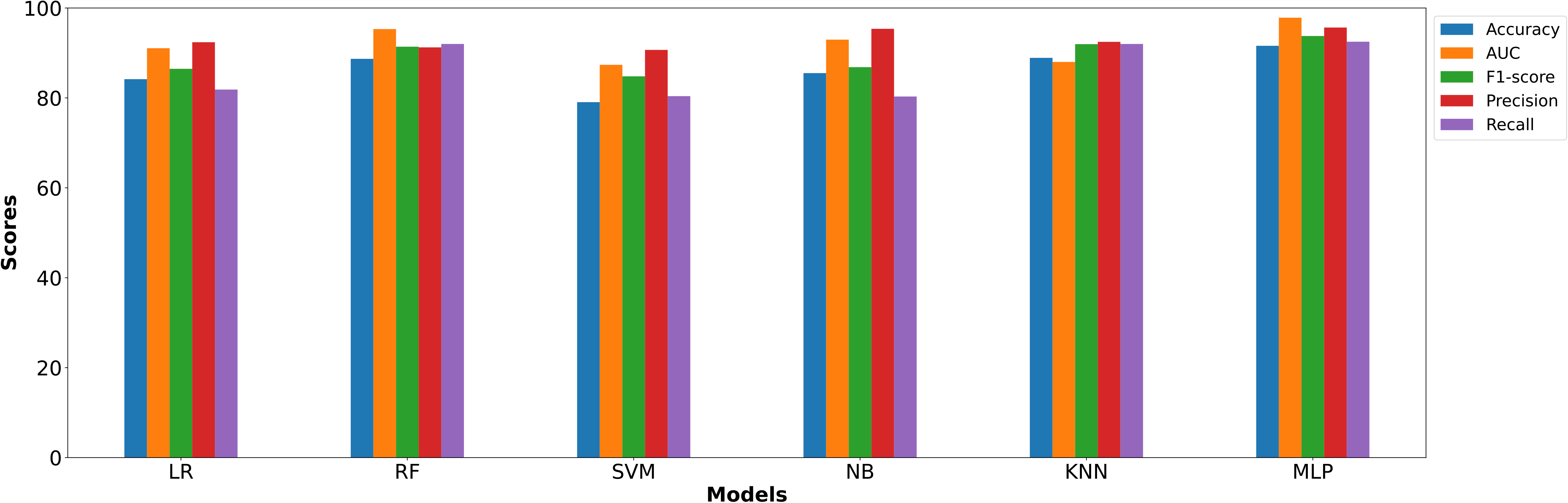
Shows a comparative analysis of the parameters of the models.

## DISCUSSION

Rice is one of the most important crops and a staple food for about half the global population. It is expected that future rice demand will increase due to the rising trend in rice consumption and the growing global population. Moreover, its production is threatened by various biotic stresses, such as bacteria, fungi, viruses, and nematodes, compromising crop yield and quality. However, rice consists of an immune system composed of PTI and ETI to counter the invasion of pathogens. The immune system is controlled by various disease-resistance proteins, such as R-proteins and PRRs. Thus, the identification of disease-resistance proteins not only helps in reducing pesticide application to rice field but also improve its yield. The development of disease resistance in rice has always been a primary goal for breeding programs, and early detection of these proteins is a major step toward this goal. Although computational methods, such as ML and DL models, can identify disease-resistance proteins, very few consider sequence-derived structural and physicochemical features of proteins. As disease-resistance proteins are crucial in rice breeding programs, their rapid and accurate prediction is needed. This work aimed to develop a model by performing a comparative analysis of six different types of classifiers, i.e., LR, RF, SVM, NB, KNN, and MLP. The best model was selected and validated based on various performance measures of these models. We identified that the DL-based MLP model was the best model over others and further studied it according to different model parameters.

By utilizing ML and DL algorithms, plant breeders can precisely identify the most advantageous traits and characteristics of plants for their specific requirements (Yoosefzadeh Najafabadi et al., 2023). Firstly, we collected a proteomics dataset, including known disease-resistance proteins called the positive dataset, and the whole proteome named the negative dataset for 13 species related to rice. Next, we performed data pre-processing steps on the negative dataset to normalize the size of both datasets for further analysis. These steps include the removal of proteins with disease-resistant domains and ‘X’ sequences. In addition, clustering was performed to reduce the redundancy in the negative dataset. Further, feature extraction and selection were done on various features of the proteins. Moreover, analyzing the performance of the six classifiers through the confusion matrix, we found that the MLP model has the highest TP values. 10-fold cross-validation on all the classifiers showed the best validation accuracy of 91.59% in the DL-based MLP model. This depicts that the MLP model is the most efficient model in predicting rice disease-resistance proteins compared to others. Furthermore, analysis of different parameters of classifiers, including AUC, F1-score, precision, and recall, showed that the MLP model outperformed other models.

In summary, the current study accurately and precisely identified disease-resistance proteins of rice and related species through the MLP model. The model is based on sequence-derived structural and physicochemical features of proteins. Thus, it predicts the disease-resistance proteins with good scores in various performance metrics. This model will help rice breeders, as well as biotechnologists, to detect disease-resistance proteins rapidly. These proteins can then be used to develop disease-resistant rice varieties in less time.

## Author Contributions

VD collected study data, performed primary analysis, and designed the working strategy; SB carried out basic data analysis and contributed to manuscript writing; performed analysis and contributed to manuscript writing; PY conceptualized and supervised the study; all authors contributed to writing the manuscript.

## ACKNOWLEDGEMENTS AND FUNDING

PY acknowledges the support from a seed grant (project number I/SEED/PY/20200037) funded by the Indian Institute of Technology, Jodhpur, India; VD and AS are thankful for the support from the Ministry of Human Resource and Development (MHRD), India fellowship.

## SUPPLEMENTAL DATA

The supplemental data is available in the online version of this article.

## DATA AVAILABILITY

All the codes are available in the Github repository (https://github.com/VedikaaDhiman/RiceDRDLM)

