## Supplemental file for "Deep learning-based method to identify disease-resistance proteins in *Oryza sativa* and relative species"

**Supplementary Tables**

**Table S1.** Overview of different databases and tools used in the study.

| **Categories** | **Description** |
| --- | --- |
| **Databases** | |
| RiceRelativesGD V4.0 database | Genomic database of phylogenetically related species of rice |
| **Tools** | |
| DRAGO 3 | Web-based tool in PRGdb database for providing a comprehensive overview of resistance genes (R-genes) in plants |
| CD-HIT | Program for clustering and comparing protein or nucleotide sequences |
| BLASTP | Tool of NCBI that compares one or more protein query sequences to a subject protein sequence or a database of protein sequences |
| protr | Package in R program, which includes a comprehensive toolkit for generating various numerical representation schemes of protein sequences |
| *Scikit-learn* | An open-source data analysis library in Python program that integrates a wide range of machine learning algorithms for supervised and unsupervised problems. |

**Table S2.** List of various descriptors computed by protr

| **Descriptor groups** | **Descriptors** |
| --- | --- |
| Amino acid composition | Dipeptide composition |
|  | Amino acid composition |
|  | Tripeptide composition |
|  | Normalized Moreau-Broto |
| Autocorrelation | Moran |
|  | Geary |
| Composition, Transition, Distribution | Composition |
|  | Transition |
|  | Distribution |
| Conjoint Triad | Conjoint Triad |
| Quasi-sequence-order | Sequence-order-coupling number |
|  | Quasi-sequence-order descriptors |
| Pseudo-amino acid composition | Type I |
|  | Type II |
| Profile-based | PSSM |
| Proteochemometric modeling | Principal components analysis (amino acid properties based) |
|  | Principal components analysis (2D and 3D molecular descriptors based) |
|  | Factor analysis (amino acid properties based) |
|  | Factor analysis (2D and 3D molecular descriptors based) |
|  | Multidimensional scaling (amino acid properties based) |
|  | Multidimensional scaling (2D and 3D molecular descriptors based) |
|  | BLOSUM and PAM matrix-derived descriptors |
